# UnigeneFinder: An automated pipeline for gene calling from transcriptome assemblies without a reference genome

**DOI:** 10.1101/2024.08.19.608648

**Authors:** Bo Xue, Karine Prado, Seung Yon Rhee, Matt Stata

**Author notes:** Corresponding authors: Matt Stata and Seung Yon Rhee.

## Abstract

For most species in nature, transcriptome data is much more readily available than genome data. Without a reference genome, however, gene calling is cumbersome and inaccurate due to the high degree of redundancy in *de novo* transcriptome assemblies. To simplify and increase the accuracy of *de novo* transcriptome assembly in the absence of a reference genome, we developed UnigeneFinder. Combining several clustering methods, UnigeneFinder substantially reduces the redundancy typical of raw transcriptome assemblies. This pipeline offers an effective solution to the problem of inflated transcript numbers, achieving a closer representation of the actual underlying genome. UnigeneFinder performs comparably or better, compared to existing tools, on plant species with varying genome complexities. UnigeneFinder is the only available transcriptome redundancy solution that fully automates the generation of primary transcript, coding region, and protein sequences, analogous to those available for high quality reference genomes. These features, coupled with the pipeline’s cross-platform implementation and focus on automation and an accessible user interface, make UnigeneFinder a useful tool for many downstream sequence-based analyses in non-model organisms lacking a reference genome, including differential gene expression analysis, accurate ortholog identification, functional enrichments, and evolutionary analyses. UnigeneFinder also runs efficiently both on high-performance computing (HPC) systems and personal computers, further reducing barriers to use.

## BACKGROUND

Despite the declining cost of genome sequencing, most species still lack a reference genome (Fig. 1). This gap poses significant challenges for biological research. RNA-seq technology provides a low-cost method of obtaining both sequence information and gene expression data. Therefore, methods for utilizing transcriptome data in the absence of a reference genome are critical for many areas of biological research. For example, in the ancient and species-rich clades Arthropoda, Embryophyta, Fungi, and Vertebrata, the number of species with RNA-seq data available is 4-11 times higher than the number of species with complete reference genomes (Figure 1). Defining the gene set of an organism from *de novo* assembled RNA-seq data thus continues to play a pivotal role in the study of non-model organisms, yet identifying the complete and accurate gene set from RNA-seq data without a reference genome remains challenging (Wang & Gribskov, 2017; Hsieh *et al*., 2019; Raghavan *et al*., 2022). *De novo* transcriptome assembly reconstructs transcripts directly from short-read sequencing data, which is inherently challenging due to factors such as alternative splicing, uneven gene expression levels, and sequencing errors (Martin & Wang, 2011; Vijay *et al*., 2013; Hölzer & Marz, 2019; Li *et al*., 2019). Consequently, assemblies are often cluttered with numerous redundant and partial transcripts, complicating nearly every type of downstream analyses such as ortholog identification, gene expression quantification, functional annotation and Gene Ontology (GO) enrichment, and phylogenetic analysis (Hsieh *et al*., 2019; Raghavan *et al*., 2022). A primary aim in non-model genomics is therefore to generate high-quality gene sets from transcriptome assemblies without having access to the reference genome.

**Figure 1:**
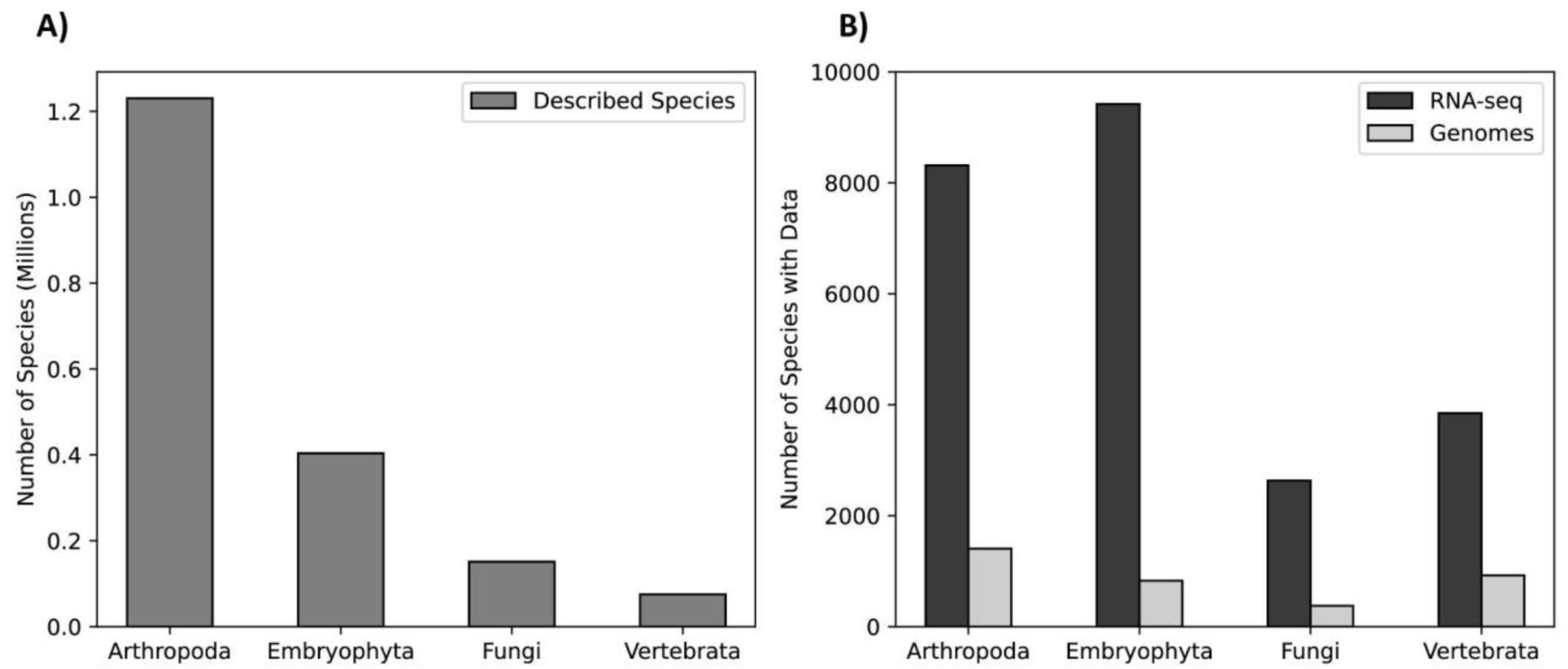
A summary of the approximate numbers of species (A) and general availability of transcriptome data versus reference genome assemblies (B) for four major clades of organisms. The gap between the number of species with RNA-seq data available and those with high-quality reference genomes remains large for major groups of organisms. In the species-rich Arthropoda, Embryophyta, Fungi, and Vertebrata clades, containing around 1.2 million, 400 thousand, 150 thousand, and 75 thousand species, respectively, there are between 4 and 11-fold more species with transcriptome data than with reference genomes. Numbers of species with RNA-seq and genomic data were obtained from NCBI, and numbers of described species were obtained from the IUCN Red List Summary Statistics (iucnredlist.org/resources/summary-statistics).

To address this challenge, we developed UnigeneFinder, a fully-automated, user-friendly ensemble pipeline that integrates diverse transcript clustering methods. UnigeneFinder converts the raw output of *de novo* transcriptome assembly software such as Trinity (Grabherr *et al*., 2011) into a set of predicted primary transcripts, coding sequences, and proteins, similar to the gene sequence data commonly available for high-quality reference genomes. UnigeneFinder achieves better precision while maintaining or improving F-scores than the individual clustering tools it combines. Unlike other available tools, it fully automates the generation of primary sequences for transcripts, coding regions, and proteins, making it suitable for diverse types of downstream analyses. This pipeline is designed to be user-friendly, cross-platform, and capable of running on personal computers, thus making it accessible to a broad range of researchers. UnigeneFinder thus aims to enable a wider range of researchers, especially those working on non-model organisms, to leverage transcriptome data for greater biological insight into species lacking a reference genome, which represent the overwhelming diversity of life.

## RESULTS

The UnigeneFinder pipeline is implemented in Python and employs several steps to derive a gene set from *de novo* assemblies of RNA-seq data without the need for a reference genome (Fig. 2). The initial input is just a transcriptome assembly in Fasta format. Trinity (Grabherr *et al*., 2011) is a commonly used *de novo* assembler program that provides a gene-to-transcript mapping file that is valuable in unigene identification. FastQ read files or BAM format read mappings can be provided for shared-read-based clustering implemented via Compacta (Razo-Mendivil *et al*., 2020). In the event that the user does not have read mappings, we implemented a second script, unigenefinder_map.py, which conducts mapping of FastQ data using hisat2 (Kim *et al*., 2019). The computational resources required for mapping reads vary depending on the quantity of read data available; larger datasets may have a long run time. UnigeneFinder then integrates the following clustering approaches with the aim of grouping transcripts by putative genomic locus: 1) the gene-to-transcript mapping produced by Trinity (Grabherr *et al*., 2011), 2) clustering based on shared read mapping using Compacta (Razo-Mendivil *et al*., 2020), and 3) sequence-based clustering using CD-HIT (Fu *et al*., 2012). The clustering information from these tools is then merged to form a network where nodes represent transcripts and edges exist between nodes if at least one clustering method grouped those transcripts together. Connected components in the network thus represent putative transcript isoforms from the same genetic locus (unigenes). Finally, from each connected component, a single transcript with the longest open reading frame (ORF) is selected as the primary transcript for the gene. The software reports a set of primary transcripts, coding sequences, and proteins for each unigene. An additional script, unigenefinder_expression.py can be utilized to generate expression data in the form of read counts and transcripts-per-million (TPM) using RSEM, which is particularly well-suited to calculating expression for transcript isoforms clustered into unigenes and provides both per-isoform and per-unigene expression information (Li & Dewey, 2011).

**Figure 2:**
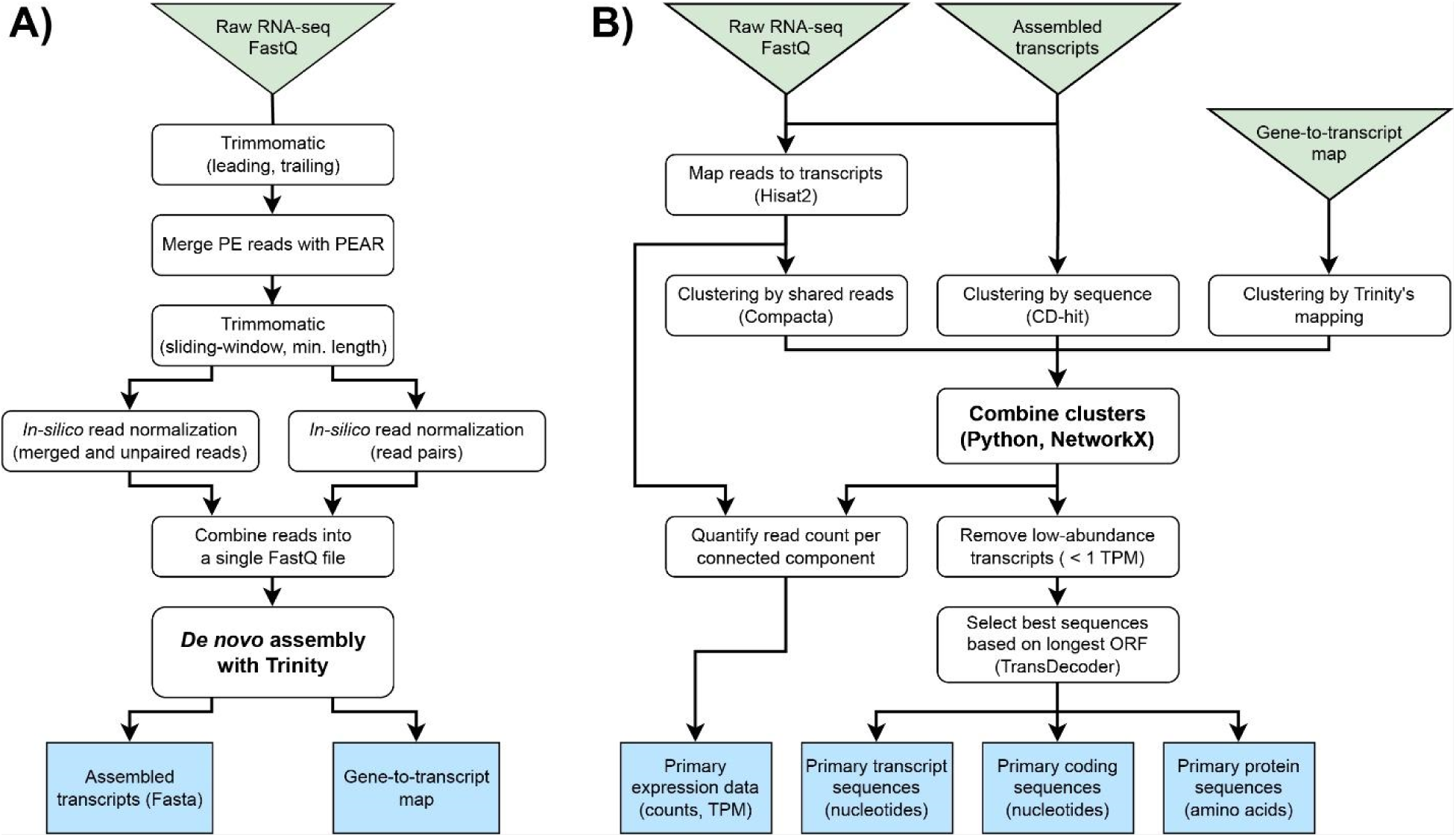
A flow diagram showing A) the recommended initial assembly process, and B) the steps implemented by the UnigeneFinder pipeline. Green triangles represent input data and blue squares represent output files. The steps in A are not included in our pipeline due to their higher computational demands and the need to be run on a high-performance compute cluster. However, scripts and example commands for these steps are included in the package.

UnigeneFinder is designed for user accessibility and efficiency. Recognizing the diverse backgrounds of researchers, we implemented a simple editable configuration file that simplifies running and customizing the pipeline, allowing users to adjust parameters to suit specific use cases. UnigeneFinder requires numerous software dependencies, both command line tools and Python libraries, the installation of which can be challenging. Therefore, we encapsulated all these dependencies in a Singularity container, which can be run with minimal difficulty on Linux, Windows, or Mac OS systems, and include instructions for each operating system (github.com/TheRheeLab/UnigeneFinder). The UnigeneFinder pipeline is thus a cross-platform solution which automates all steps from a raw assembly produced by Trinity or similar assembly software to a set of putative primary transcripts, coding sequences (CDS), and protein sequences, plus expression information. The output thus mirrors the organization of data typically associated with high-quality reference genomes, facilitating downstream analyses such as quantification of gene expression, functional annotation, gene function enrichment, ortholog identification, and evolutionary analysis.

We next benchmarked UnigeneFinder against the individual clustering tools that were integrated into our pipeline by deploying each tool on *de novo* transcriptome assemblies of four plant species with high-quality chromosome-scale reference genomes (Table 1). We selected these species based on their genome size and complexity to represent a wide array of use cases. *Arabidopsis thaliana* is a eudicot with a relatively compact and simple genome, with an assembly size of 120 Mb, whose most recent genome duplication event occurred in the common ancestor of the Brassicaceae family (The Arabidopsis Genome Initiative, 2000; Barker *et al*., 2009; Koenig & Weigel, 2015). *Solanum lycopersicum*, another eudicot with a genome assembly size of 802 Mb, is a member of the Solanaceae family, whose shared ancestor underwent a genome triplication (Sato *et al*., 2012; Barchi *et al*., 2019). *Brachypodium distachyon* has a 270 Mb genome assembly size and is a member of the grass family Poaceae, which underwent three ancient genome duplication events (McKain *et al*., 2016). *Zea mays* has a genome assembly size of 2.18 Gb and shares the same three grass family duplications plus an additional recent duplication occurring 5 to 12 million years ago (Schnable *et al*., 2009). Together, these four species represent a broad range of genome complexities with which to test the performance of UnigeneFinder.

**Table 1:** Data sources and genome size of species used for benchmarking UnigeneFinder. RNA-seq data used was current for all NCBI BioProjects as of April 17, 2024.

| Species | Genome Version | Assembly Size (Gb) | Number of Scaffolds | Min. Scaffolds Containing Half the Assembly | Protein-Coding Genes | NCBI BioProject(s) for RNA-seq Data Used |
| --- | --- | --- | --- | --- | --- | --- |
| *Arabidopsis thaliana* | Araport11 (Phytozome ID: 447) | 0.12 | 7 | 5 | 27,655 | PRJNA910862 |
| *Solanum lycopersicum* | ITAG5.0 (Phytozome ID: 796) | 0.802 | 13 | 6 | 36,648 | PRJNA339202<br>PRJNA645803<br>PRJNA975110 |
| *Brachypodium distachyon* | 3.2 (Phytozome ID: 556) | 0.27 | 10 | 3 | 32,439 | PRJNA651067 |
| *Zea mays* | B73 5.0.55 (Phytozome ID: 833) | 2.182 | 685 | 5 | 39,756 | PRJNA918441 |

Benchmark results demonstrate that our approach substantially reduces the number of transcripts. The initial Trinity assembly of *Arabidopsis thaliana* with no post-processing yielded 117,928 assembled transcripts for only 19,366 genes with expression levels above 1 TPM in the RNA-seq samples used. Similarly, *Brachypodium distachyon*, *Solanum lycopersicum*, and *Zea mays* produced initial assemblies with 172,367, 184,342, and 400,690 transcripts, respectively, compared to their significantly lower counts of 22,000-25,000 expressed genes in these species (Tables 1 and 2). These numbers represent a 6-16 fold increase due to redundant isoforms. Sequence inflation of this nature is typical of raw *de novo* transcriptome assemblies, both provided by sequencing companies such as NovoGene and generated in house. Processing these raw assemblies with UnigeneFinder resulted in a number of transcripts much closer to the number of expected genes (Fig. 3). Our approach of integrating these different clustering methods also achieves better precision than any individual tool alone, with only a slight loss in recall, resulting in an improved or near-identical F-score relative to existing tools, for all four test assemblies (Fig. 3, Table 2). For *Arabidopsis thaliana*, with the simplest genome in the test set, UnigeneFinder achieved near identical precision compared to the Trinity clustering alone, and near identical F-Score compared with Compacta, with a difference of under 1% in each. For the three other species with greater genome complexity, UnigeneFinder achieved the best precision and F-score, with margins of improvement in precision of 2-4% and in F-scores of 1-2% (Table 2). Table 3 reports run time, CPU time, memory, and CPU use for both a personal laptop and high-performance computing (HPC) cluster node. For all four test assemblies, run times were under one hour and thus not prohibitively long even on a personal computer, indicating that HPC access is not a requirement for running this pipeline. UnigeneFinder thus matches or outperforms the individual methods that it integrates, and couples this performance with an accessible design and automated output of refined primary transcript, coding region, and protein sequence data, and because UnigeneFInder is cross-platform and can run on a typical personal computer without requiring access to HPC, the barriers to use are minimal.

**Figure 3:**
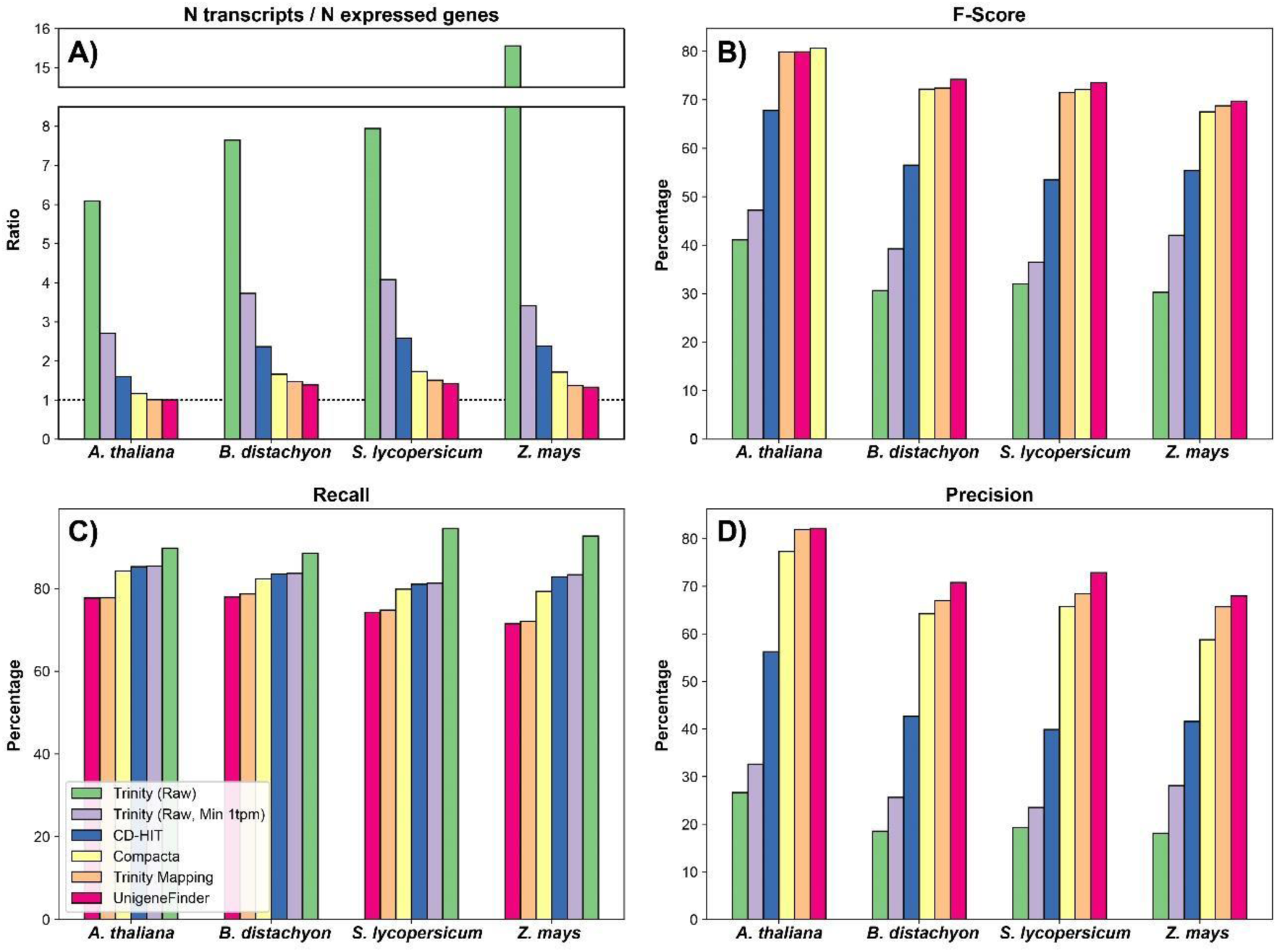
Performance comparison of UnigeneFinder and component clustering algorithms on four test assemblies of varying genomic complexity. The ratio of the number of transcripts to expressed genes (A), F-score (B), recall (C), and precision (D) are presented for *Arabidopsis thaliana* (Brassicaceae), *Brachypodium distachyon* (Poaceae), *Solanum lycopersicum* (Solanaceae), and *Zea mays* (Poaceae). Definitions of precision and recall are given in Methods. The dashed line in A denotes a one-to-one ratio.

**Table 2:** Precision, recall, F-score, and number of assembled transcripts with *de novo* assemblies of four species, comparing raw assemblies and assemblies processed with UnigeneFinder as well as the individual clustering methods. Because minimum expression filtering is included in UnigeneFinder, the component methods (Trinity’s gene-to-transcript mapping, Compacta, and CD-HIT) are all presented here with transcripts < 1 TPM removed as well.

|  |  | Method |  |  |  |  |  |
| --- | --- | --- | --- | --- | --- | --- | --- |
|  |  | Trinity (Raw) | Trinity (Raw, Min 1tpm) | Trinity Mapping | Compacta | CD-HIT | Unigene Finder |
| *Arabidopsis thaliana* | Transcripts | 117,928 | 52,499 | 19,540 | 22,563 | 30,907 | 19,471 |
|  | Recall | 89.79 | 85.39 | 77.77 | 84.24 | 85.30 | 77.70 |
|  | Precision | 26.66 | 32.61 | 81.94 | 77.31 | 56.24 | 82.15 |
|  | F-Score | 41.11 | 47.20 | 79.80 | 80.63 | 67.79 | 79.86 |
| *Brachypodium distachyon* | Transcripts | 172,367 | 84,078 | 33,151 | 37,463 | 53,291 | 31,262 |
|  | Recall | 88.57 | 83.68 | 78.70 | 82.39 | 83.53 | 77.96 |
|  | Precision | 18.52 | 25.63 | 67.02 | 64.22 | 42.68 | 70.79 |
|  | F-Score | 30.63 | 39.24 | 72.39 | 72.18 | 56.49 | 74.20 |
| *Solanum lycopersicum* | Transcripts | 184,342 | 94,694 | 34,962 | 40,131 | 59,946 | 32,926 |
|  | Recall | 94.55 | 81.30 | 74.80 | 79.87 | 81.06 | 74.22 |
|  | Precision | 19.29 | 23.52 | 68.43 | 65.76 | 39.90 | 72.84 |
|  | F-Score | 32.04 | 36.48 | 71.47 | 72.13 | 53.48 | 73.52 |
| *Zea mays* | Transcripts | 400,690 | 87,865 | 35,429 | 44,114 | 61,193 | 34,155 |
|  | Recall | 92.70 | 83.36 | 72.07 | 79.31 | 82.83 | 71.50 |
|  | Precision | 18.09 | 28.11 | 65.70 | 58.77 | 41.59 | 67.97 |
|  | F-Score | 30.27 | 42.04 | 68.74 | 67.51 | 55.38 | 69.69 |

**Table 3:** Runtime and computational resources used by UnigeneFinder for all four test assemblies, on a personal laptop computer (Microsoft Surface Book 2, Intel Core i7-8650U CPU with 1.90GHz base frequency, four cores, eight logical threads, 16GB RAM, running Windows 10) and an HPC cluster node (Intel Xeon Gold 6148 CPU with 2.40GHz base frequency, 40 cores, 80 logical threads, 202 GB RAM, running Linux 3.10.0).

|  | Laptop Computer |  |  |  |
| --- | --- | --- | --- | --- |
|  | Raw Assembly Size | Run Time (mm:ss) | CPU Time (h:mm:ss) | Peak Memory Use (gigabytes) |
| *Arabidopsis thaliana* | 117,928 | 16:22 | 0:39:54 | 0.518 |
| *Brachypodium distachyon* | 172,367 | 28:20 | 0:57:31 | 0.757 |
| *Solanum lycopersicum* | 184,342 | 34:40 | 1:08:26 | 0.832 |
| *Zea mays* | 400,690 | 47:14 | 2:29:48 | 1.267 |
|  | HPC Node |  |  |  |
|  | Raw Assembly Size | Run Time (minutes) | CPU Time (minutes) | Peak Memory Use (gigabytes) |
| *Arabidopsis thaliana* | 117,928 | 18:24 | 0:34:16 | 0.491 |
| *Brachypodium distachyon* | 172,367 | 34:27 | 0:49:04 | 0.709 |
| *Solanum lycopersicum* | 184,342 | 39:01 | 0:59:48 | 0.819 |
| *Zea mays* | 400,690 | 58:14 | 2:03:20 | 1.244 |

## DISCUSSION

The advancement of genome sequencing technologies has been both rapid and transformative, propelling the field of genomics into a new era of rapid discovery. Since the completion of the first eukaryotic genomes, sequencing costs have plummeted, making genome sequencing more accessible than ever before (Mardis, 2017). Despite these technological strides and cost reductions, the coverage of reference genomes across the tree of life remains remarkably sparse. Most species across many diverse and ecologically, evolutionarily, and agronomically significant clades lack a reference genome. This scarcity represents a fundamental barrier to progress in biological research, including functional biology, conservation biology, and evolutionary studies. In plants, the situation is particularly acute. Despite their critical role in ecosystems and agriculture, comprehensive genomic resources are available for only a fraction of the estimated 391,000 plant species (Cheng *et al*., 2018). For numerous diverse clades, such as the Asparagales, Asterales, Gentianales, and Polypodiales, the availability of reference genomes is particularly low despite their ecological significance (Marks *et al*., 2021). This scarcity of reference genomes hinders our ability to address fundamental questions about plant diversity, adaptation, and resilience. As we strive to catalog the Earth’s biodiversity and gain greater insight into adaptations that provide resilience for diverse organisms, there is a great need both for a concerted effort to increase genome sequencing coverage and for methods to leverage insight from non-model species which lack a reference genome.

*De novo* transcriptome assembly from short-read RNA-seq data has long been a crucial approach in the study of non-model organisms, facilitating the simultaneous acquisition of gene sequences and expression data without the prerequisite of a reference genome. This can be achieved at a significantly lower cost than assembling an entire genome, enabling researchers to investigate the genetic makeup and functional dynamics of a wide array of non-model species, thus broadening our understanding of biological complexity and evolution. However, despite its importance, *de novo* transcriptome assembly is still fraught with challenges. The intrinsic variability of several orders of magnitude in gene expression levels and differences in transcriptome patterns across different tissues and developmental stages, combined with the occurrence of alternative splicing, adds substantial complexity to the assembly process (Martin & Wang, 2011; Grabherr et al., 2011). These factors often result in the production of assembled transcripts that far exceed the actual number of genes within the organism’s genome, complicating downstream analyses such as functional annotation, differential expression, and evolutionary comparisons (Vijay *et al*., 2013; Haas *et al*., 2013).

In this study, the disparity between the number of transcripts obtained via *de novo* RNA-seq assembly and the estimated number of expressed genes obtained from the reference genomes highlights this challenge, with Trinity producing between 6 and 16 times more transcripts than there are expressed genes in each test case. Such discrepancies underscore the difficulty of *de novo* assembly and the potential for redundancy in assembled transcripts (Haas et al., 2013; Hölzer & Marz, 2019). These excess transcripts, many of which represent fragmented or otherwise incomplete assemblies, can interfere with almost every type of downstream analysis and obscure true biological signals (Vijay et al., 2013; Hölzer & Marz, 2019). Ortholog prediction, phylogenomic studies, gene ontology (GO) enrichment, and gene expression analysis are all prime examples of analyses whose accuracy suffers due to redundant assemblies. For example, phylogenomic inference typically uses single or low copy orthologs and requires the exclusion of paralogs. Therefore, the presence of numerous transcripts for a single locus can complicate the differentiation between orthologous and paralogous sequences, greatly reducing the set of usable genes and potentially leading to inaccurate evolutionary interpretations (Altenhoff *et al*., 2019). Similarly, gene expression analysis is adversely affected by the presence of redundant transcripts, since when reads cannot be unambiguously mapped to one isoform or another they are typically randomly assigned and thus dilute read counts, resulting in an underestimation of total expression level per gene (Martin & Wang, 2011). This can be particularly confounding in comparisons across different assemblies, where the number of isoforms in each (and thus the degree to which expression is affected) may vary. At the same time, the presence of multiple isoforms of a gene that responds to experimental conditions can artificially inflate the number of differentially expressed genes, obscuring the true transcriptomic landscape of the organism under study, and affecting downstream analyses such as identification of enriched GO terms among responsive genes. Thus, while *de novo* transcriptome assembly opens new avenues for genetic and functional studies of non-model organisms, it also necessitates tools for removing redundancy and identifying primary isoforms for each gene without a genome sequence or knowledge of the true genetic loci.

To better leverage transcriptomic data for biological insights in non-model species, we developed UnigeneFinder, an automated pipeline designed to reduce the number of redundant genes in *de novo* RNA-seq gene assemblies. By integrating diverse clustering methods - Trinity’s gene-to-transcript mapping produced during De Brujin graph-based assembly (Grabherr *et al*., 2011), Compacta’s clustering based on shared read mapping (Razo-Mendivil *et al*., 2020), and CD-HIT’s sequence similarity-based clustering (Fu *et al*., 2012) - the UnigeneFinder pipeline optimizes the accuracy of transcript assembly, achieving the highest performance in precision and the least inflation in gene numbers across four test assemblies of varied genomic complexities. The resulting ratios of assembled genes to the expected number of expressed genes from the genome—1.01X for Arabidopsis, 1.39X for Brachypodium, 1.42X for Solanum, and 1.33X for Maize—are much closer to these species underlying genomes than the original assemblies, which exhibited 6-16 times more assembled genes than expected expressed genes. By automating these complex processes and producing accurate and minimally redundant gene sequence data, UnigeneFinder serves to both increase the utility of transcriptome data in the absence of a reference genome. UnigeneFinder is bundled with all its software dependencies in a cross-platform and user-friendly manner and does not require HPC access, thus lowering barriers to entry for conducting genomics research on non-model organisms.

## CONCLUSION

UnigeneFinder represents a useful tool for leveraging more from transcriptome data from non-model species lacking a reference genome. This software pipeline provides a robust solution for refining raw transcriptomic data into a curated set of gene sequences coupled with expression data. Its integration of multiple clustering methods enhances precision without substantially sacrificing recall. The focus on a user-friendly cross-platform design and availability as an open-source pipeline ensures that it is accessible to a broad spectrum of users. As sequencing costs continue to decrease, resulting in rapidly increasing amounts of raw data, tools such as UnigeneFinder will play an important role in generating new insights from transcriptome data of diverse and understudied branches of the tree of life, contributing to our understanding of biological complexity and innovation.

## METHODS

### Pipeline structure

A pipeline of existing software tools was assembled with the aim of accurately clustering *de novo* transcriptome assemblies down to the best isoforms representing unique loci in the genome (Fig. 2). Three distinct clustering methods were used: the gene-to-transcript mapping produced by Trinity (Grabherr *et al*., 2011), sequence similarity-based clustering using CD-HIT (Fu *et al*., 2012), and shared-read-based clustering using Compacta (Davidson & Oshlack, 2014; Razo-Mendivil *et al*., 2020). These different clustering methods were then integrated using a Python script which builds a network representation of the clusters using the networkx Python package (networkx.org). Nodes represent transcripts and edges exist between any two transcripts which were grouped together by one or more of these clustering approaches. Best genes were selected per cluster based on the longest open reading frame, predicted using TransDecoder (github.com/TransDecoder/TransDecoder). The entire process was automated with Python and encapsulated in a Singularity container (Kurtzer et al., 2017) in order to include all software dependencies. The pipeline is therefore cross-platform, and can be run natively on Linux systems or via the Windows Subsystem for Linux (docs.microsoft.com/en-us/windows/wsl/) on a PC or using Docker for Mac (docs.docker.com/docker-for-mac/) in Mac OS.

### Benchmarking

Paired-end RNA-seq data for four species were downloaded from the NCBI SRA (Table 1). Reads were trimmed based on quality using Trimmomatic (Bolger *et al*., 2014) with the parameters ILLUMINACLIP:Adapters.fa:5:5:5 LEADING:35 TRAILING:35 MINLEN:120 to remove any sequencing adapters and trim low-quality bases from either end. Next, PEAR was used to merge paired-end reads into single longer reads with a p-value parameter of 0.001 (Zhang *et al*., 2014). Assembled and unassembled reads were then trimmed again using Trimmomatic with parameters SLIDINGWINDOW:5:30 MINLEN:100 to remove reads with low quality internal sequences and retain only reads with a minimum length of 100 bp. *In silico* read normalization was conducted using the Perl script included with Trinity, with min and max coverage parameters of 2 and 100, respectively. Trinity was used with default parameters to assemble the transcriptome of each species using these normalized reads.

For each of these four test assemblies, the number of assembled genes, recall, and precision were evaluated. Because transcriptome data necessarily only capture a subset of genes expressed in the sampled tissues, we limited calculations of precision and recall to genes with at least 1 transcript per million (TPM) expression, which is a widely used minimum threshold in gene expression studies. Expression levels were quantified using hisat2 to map RNA-seq read data onto the primary coding sequences of each reference genome, followed by SAMTools for generating sorted read pile-ups in BAM format (Kim *et al*., 2019; Danecek *et al*., 2021). From these data, TPM values were calculated using Python. TPM values were calculated in the same way for the assembled transcripts for use in filtering out low-abundance transcripts, with this step implemented for the UnigeneFinder pipeline and each component step in order to achieve an unbiased comparison of methods.

Precision, recall, and F-scores were calculated in the standard manner: with A as true positives, B as false positives, and C as false negatives:

**Precision** = A / (A + B), and **Recall** = A / (A + C)

F-scores were calculated as the harmonic mean of precision and recall, or:

**F-score** = 2[(Precision * Recall) / (Precision + Recall)]

True positives, false positives, and false negatives were defined based on nucleotide BLAST against coding sequences for each gene in the reference genome. We defined true positives (A) as the best assembled transcript mapping to a given gene via BLAST, while all subsequent transcripts mapping to a gene were considered false positives (B). False negatives (B) were defined as genes in the genome which met the minimum 1 TPM expression requirement but lacked an associated assembled transcript. UnigeneFinder was tested on a laptop equipped with an Intel Core i7-8650U CPU with 1.90GHz base frequency, four cores, eight logical threads, 16GB RAM, running Windows 10, as well as on an HPC cluster node with two Intel Xeon Gold 6148 CPUs each 20 cores and 40 logical threads, with a 2.40GHz base frequency, and 202GB RAM, running Linux 3.10.0. Run time, CPU time, and CPU and memory use for both test systems were recorded.

### Pipeline and data availability

All RNA-seq data used for benchmarking UnigeneFinder are from public bioprojects on the NCBI Sequence Read Archive (ncbi.nlm.nih.gov/sra), with project IDs listed in Table 1. The UnigeneFinder pipeline is available via GitHub (github.com/TheRheeLab/UnigeneFinder).

## Supplemental Tables

**Table S1:** Version information for all software tools used.

| Software | Version | Reference |
| --- | --- | --- |
| CD-HIT | 4.8.1 | Fu et al., 2012 |
| Compacta | 1.02 | Razo-Mendivil et al., 2020 |
| HISAT2 | 2.2.1 | Kim et al., 2019 |
| SamTools | 1.13 | Danecek et al., 2021 |
| SRA-Toolkit | 3.0.9 | trace.ncbi.nlm.nih.gov/Traces/sra/sra.cgi?view=software |
| Trimmomatic | 0.39 | Bolger et al., 2014 |
| Trinity | 2.15.1 | Grabherr et al., 2011; Haas et al., 2013 |
| BLAST+ | 2.14.1 | Camacho et al., 2009 |
| TransDecoder | 5.7.1 | github.com/TransDecoder/TransDecoder |

